# Life History Optimization and the Macroevolution of Mammal Body Size

**DOI:** 10.1101/2025.10.22.684073

**Authors:** Jack da Silva

## Abstract

Although body size is linked to nearly every aspect of an organism’s biology, its evolution remains poorly understood. Here, an analytical life history model of optimal size is tested for the first time using unprecedented data on mammal life history and phylogeny and shown to explain several well-defined macroevolutionary patterns. The evolution of body size is explained in large part by life history optimization with respect to adult mortality under metabolic constraints on productivity. The model also explains plausible effects of climate change, diet, feeding mode, cursoriality, aquatic living, powered flight, and island endemicity on the evolution of body size. This study furthers our understanding of the relationship between micro- and macro-evolution.

## Introduction

We do not fully understand the evolution of body size ^1-3^. This is remarkable considering that body size is integral to an organism’s physiology, life history, ecology, and behaviour, as well as a species’ temporal and geographical distribution and species diversity ^2-8^. Two approaches have been used to remedy this. First, life history theory may be used to predict the size that maximizes fitness given ecological circumstances and anatomical, physiological and genetic constraints ^9,10^. This microevolution approach has been successful in that it provides a clear logical basis for gradual evolutionary change within populations over short time scales, but has generally not been verified for body size because of the difficulty of estimating model parameters ^1,11-14^. In addition, this approach does not usually identify the external forces, such as aspects of environmental change, that have imposed selection on body size. A second approach is to correlate variation in body size among species or higher taxa over time and space with external factors as a means of identifying the forces changing body size ^3,15^. This macroevolution approach has identified many plausible factors that are potentially important, but these hypotheses are inherently difficult to test because of their time scales. In addition, these hypotheses do not specify the evolutionary processes in play except in very general terms, such as whether they are adaptive, stochastic, gradual, punctuational, anagenic, or cladogenic ^16-24^. Neither approach on its own provides a satisfactory picture of the evolution of body size. Here, a life history model of optimal body size is developed and tested in mammals and used to predict well-defined macroevolutionary patterns. This work establishes conceptual and empirical bridges between micro- and macro-evolution ^25^, providing a more complete understanding of body size evolution.

The class Mammalia provides a useful source of data with which to describe and test both micro- and macro-evolutionary processes and patterns of body size evolution (Figure 1). Mammals originated in the Mesozoic Era ^26^ and underwent an explosive radiation after the Cretaceous–Paleogene mass extinction, ∼66 millions years ago, exhibiting a net increase in size throughout the Cenozoic Era ^18,21,27,28^. Adult body masses of the ∼6700 species of extant mammal ^29^ span eight orders of magnitude (84 million-fold difference), from the 1.77 g Sulawesi tiny shrew (*Crocidura levicula*) to the 149,000 kg blue whale (*Balaenoptera musculus*) ^30^, the largest animal that has ever lived. In addition, substantial information is available on the phylogeny, ecology, and life history of extant and fossil mammals ^30-34^.

**Figure 1.**
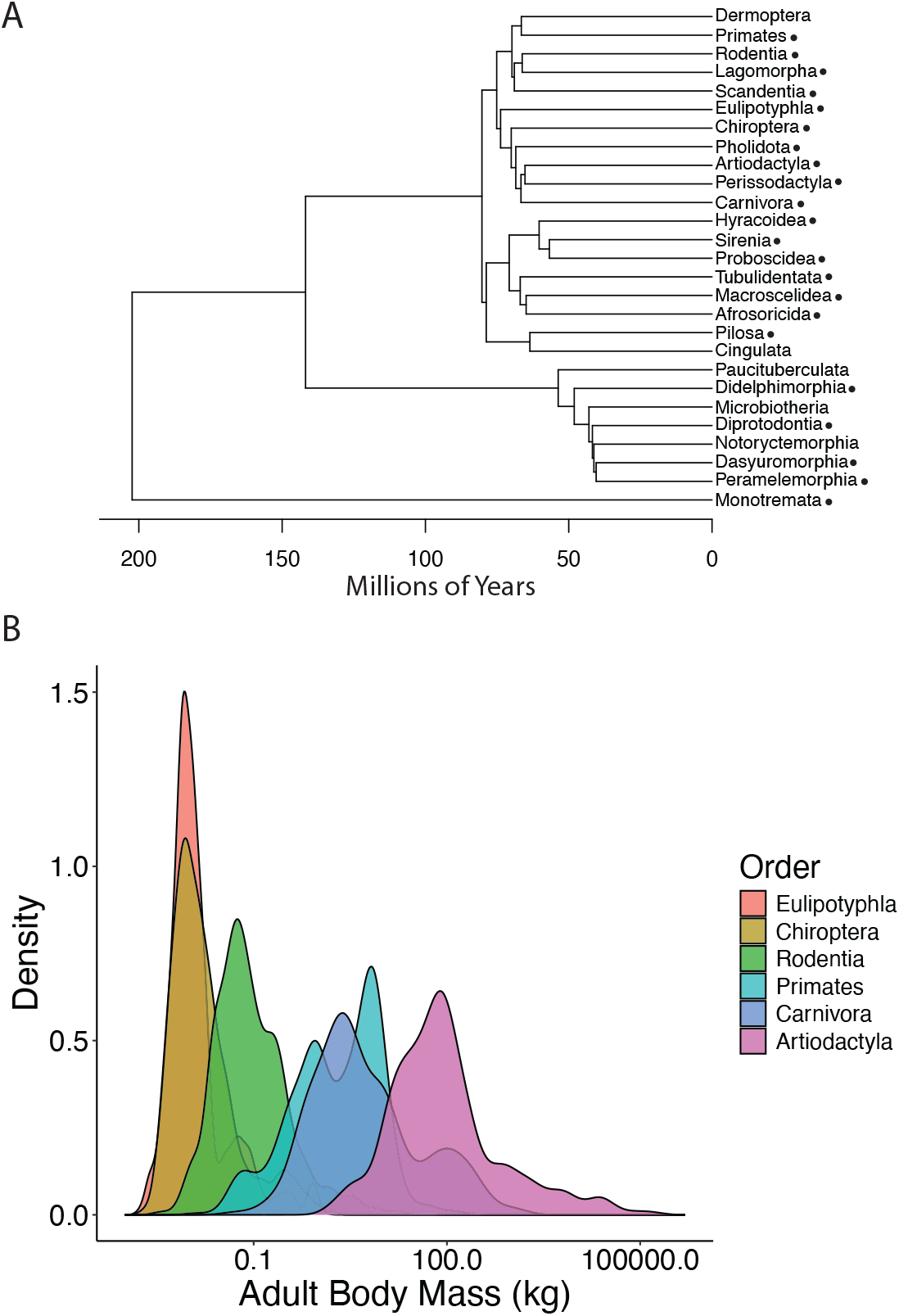
Data sources. (A) Phylogeny of the 27 orders of extant mammals from the species-level phylogeny used in this study ^31^. Dots indicate the 22 orders for which the necessary data are available for this study. (B) Density distributions of adult body mass for highly sampled mammal orders (n > 200 species each; 5000 species in total) in the COMBINE database ^30^, spanning the range of mammal body sizes.

The rate of increase in body size during the Cenozoic varies over time and among lineages and is generally higher for heavier lineages ^16,18,19,21,27,28,35,36^. Larger size is also associated with higher speciation and lower extinction rates in some clades ^17,24^. Various drivers of these changes have been proposed, including new niches and climate and associated vegetation changes ^18,24,28,35,37-41^. For example, adaptive shifts related to diet or mode of living, such as powered flight and aquatic living, are argued to select for changes in body size ^23,36,37,42-46^. Biogeography may also be important ^28,36,40^. For example, large mainland species tend to become smaller on islands, whereas small mainland species tend to become larger on islands ^36,47-50^.

Some authors have expressed doubts that life history optimization, or microevolutionary processes more generally, can explain macroevolutionary patterns of mammal body size ^16-18,23,24,51,52^. This stems mainly from evidence of punctuated evolution, species sorting, or cladogenesis. However, recent analyses of the evolution of body size in phylogenies of extant mammals generally find that its tempo and mode are consistent with Darwinian gradualism ^19-21^. Life history theory may provide insights into the details of the microevolutionary processes involved. In species with determinate growth, such as mammals, individuals typically grow until they begin reproduction. Life history theory predicts that the optimal adult size is reached when switching from growth to reproduction maximises fitness ^11-14^. However, for such models to be testable they must be simple enough to be analytically tractable and make predictions based on readily measured variables and estimable parameters. The analytical nature of such a model will also provide an intuitive understanding of body size evolution. As a recent counter example, a complex numerical computational model of life history evolution developed to predict optimal body size turns out to be difficult to test because of its numerous parameters ^1^. Here, an analytical model of lifetime reproductive effort based on metabolic scaling ^53^ is used to predict optimal body size. The model is tested by utilizing an unprecedented amount of data from a coalesced database of mammal life history and ecological data ^30^ and a recent fossil-calibrated, time-scaled molecular phylogeny ^31^ containing most extant species. The model accurately predicts adult body mass across mammals. In addition, the model explains well-documented macroevolutionary patterns that suggest some of the important external selective forces that have acted on body size: climate and associated vegetation changes, diet, feeding mode, cursoriality, aquatic living, powered flight, and island endemicity.

### The model

The model is based on the species-specific production rate, *P*(*w*), as a power (allometric) function of body size, *w*: *P*(*w*) = *aw*^*b*^, where *a*, the scaling coefficient, and *b*, the scaling exponent, are model parameters. The production rate is the difference between assimilation and respiration rates, and in the simplest, and thus analytically tractable, model it is the amount of energy allocated to growth or reproduction. Then, measuring fitness as lifetime reproductive success (see Supplementary Information), growth should stop, and reproduction begin, when the rate of increase of production with respect to body size is equal to the instantaneous adult mortality rate ^11-14^: 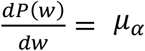, and therefore, 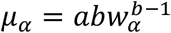, where *α* is the age of maturity. This model is difficult to test because the parameter *a* varies considerably among taxonomic and functional groups in a manner that is not fully understood, but which likely reflects external selective factors ^6,54,55^. However, in analyzing variation in lifetime reproductive effort (*LRE*) in mammals and lizards, Charnov, et al. ^53^ showed that *LRE* may be written in terms of the allometric production rate functions defining optimal size, providing a way to predict optimal size at maturity without estimating *a*. They did this by equating the reproduction rate with the production rate (assumed to be equal in the optimization model). The reproduction rate is the product of the rate of fecundity, *m*, and offspring mass at independence (mean mass of an offspring reaching independence through weaning or death), *w*_*i*_. As such, *LRE* is the reproduction rate relative to adult mass for the duration of the reproductive lifespan (life expectancy at maturity, *E*_*α*_): 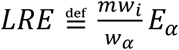. *LRE* may then be written in terms of the allometric production functions of the optimization model, using 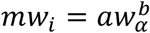 and 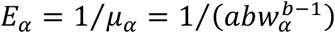, to give

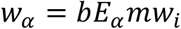

That is, optimal size at maturity is equal to the scaled lifetime production of independent offspring mass. This model suggests that an organism should grow until investment in somatic tissue (body mass) can be recouped by reproduction over its reproductive lifespan ^56^. Values for the demographic and life history variables in the model and the production rate scaling exponent are available or estimable for many species of mammal. Details and tests of the model’s assumptions are provided in Supplementary Information.

However, two important assumptions suggest that the model as presented is incomplete. First, the reproduction rate, *R* = *mw*_*i*_, based on the production of offspring mass at independence, is likely to underestimate actual reproductive effort because indirect reproduction costs (energetic costs other than the energy content of offspring) account for the majority of reproduction costs in mammals ^57^. Unfortunately, the proportions of these costs have been estimated for very few species. On the other hand, the simplifying assumption in the model that in adulthood all energy is allocated to reproduction, made for analytical tractability, overestimates the proportion of the total production rate directed to reproduction because some production must be allocated to somatic maintenance ^58^. Again, this proportion is unknown. To correct the model, it may be assumed that the true reproduction rate lies between *R* and the maximum production rate, *P*_*max*_. *P*_*max*_ may be estimated as the maximum growth rate, *k*_*max*_ (see Methods). As expected, using *k*_*max*_ in place of *R* in the model overestimates adult body mass (Supplementary Information). Therefore, the true reproduction rate is assumed to be some proportion of *k*_*max*_. Using an adjusted reproduction rate of *R*_*adj*_ = 0.35*k*_*max*_ in the model (*w*_*α*_ = *bE*_*α*_*R*_*adj*_) gives a good match between predicted and observed adult body mass, suggesting that the true reproduction rate is ∼1/3 of the maximum production rate.

### Testing the model

The model was tested by comparing predicted and observed adult body masses. The necessary demographic data (*E*_*α*_) could be estimated directly from population data for 49 species in eight orders (“Demography” dataset; see Methods). Forty-one of these species, in eight orders, also have adult, neonate, and weaning body masses and the age at weaning in the COMBINE database ^30^ (see below), allowing estimation of optimal adult mass and its comparison to observed adult mass. The model gives an excellent match to observed body mass (PGLS log-log regression; *r*^2^ = 0.94), with with an intercept (−0.02) not significantly different from 0 on a log_10_ scale (*t* = −0.3215, *P* = 0.7495) and a slope (1.00) not significantly different from 1 (*t* = 0.1156, *df* = 39, *P* = 0.9086) (Figure 2A).

**Figure 2.**
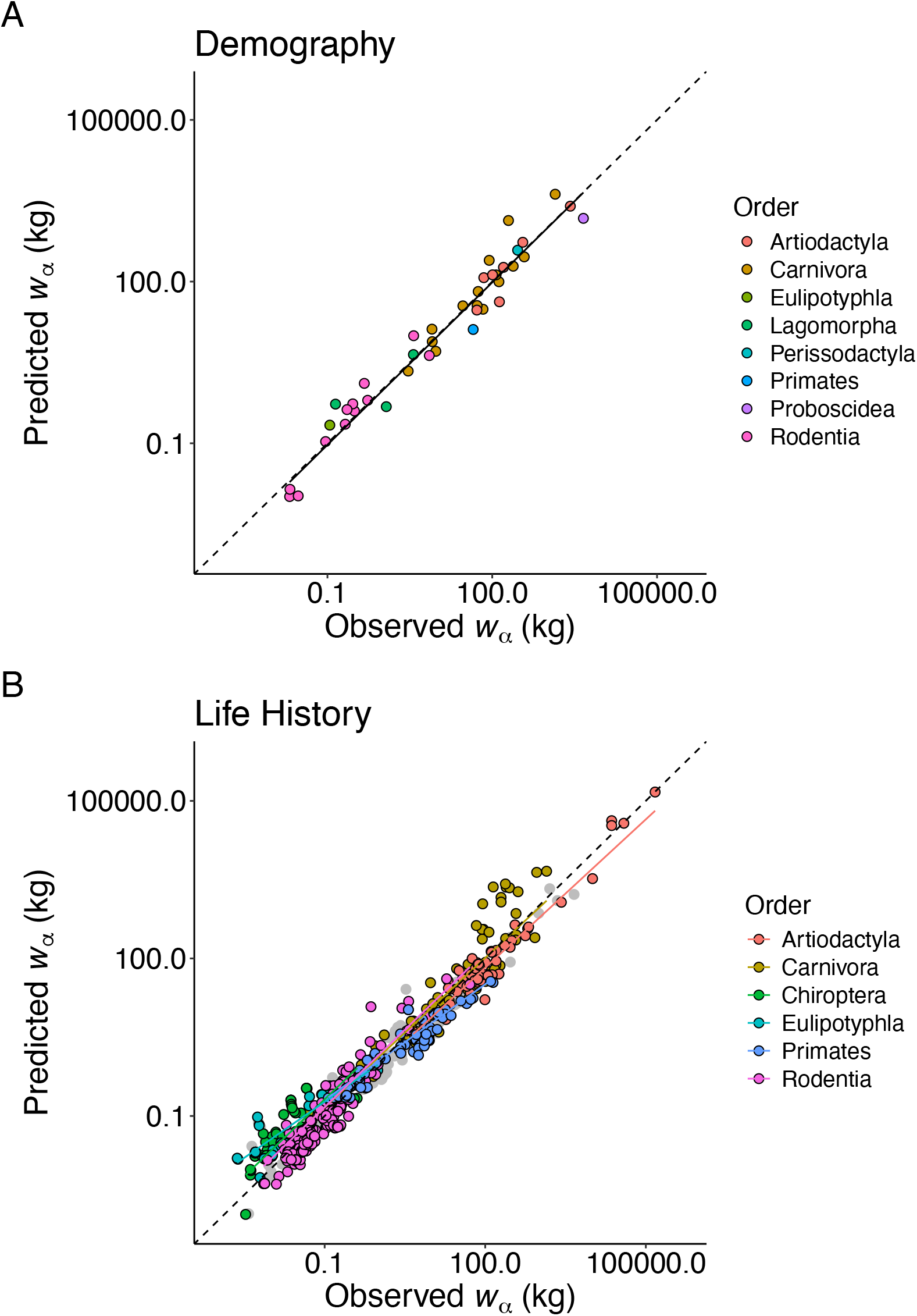
Testing the life history model. (A) PGLS log-log regression of predicted (optimal) adult body mass on observed mass using the Demography dataset (41 species in 8 orders): λ = 0.93, κ = 1.09, δ = 2.92; r^2^ = 0.94, F_1,39_ = 609.1, P < 2.20 × 10^−16^; y = −0.02 + 1.00x. (B) PGLS log-log analysis of covariance (ancova) of the effect of order on predicted adult mass, with observed mass as a covariate, using highly sampled orders in the Life History dataset (571 species in the phylogeny): λ = 1.00, κ = 0.94, δ = 2.88; r^2^ = 0.83, F_11,559_ = 241.5, P < 2.20 × 10^−16^. Grey datapoints are from species not in the analysed orders or not in the phylogeny. Dashed lines indicate y = x.

The model was also tested using a much larger dataset derived from the COMBINE database ^30^. The COMBINE database contains data on the age of female first reproduction, the age at weaning, and adult, neonate, and weaning body masses for 794 species in 22 of the 27 extant orders, comprising the “Life History” dataset. Life expectancy at maturity (*E*_*α*_), which is difficult to estimate and thus rarely reported, is not available in the database. Therefore, *E*_*α*_was estimated from its strong relationship with the age at first female reproduction (*α*), estimated using the Demography dataset (see Methods). In the life history dataset, adult mass ranges over more than seven orders of magnitude, from 2.33 g for the Sri Lankan shrew (*Suncus fellowesgordoni*) to 1.49 × 10^8^ g for the blue whale (*Balaenoptera musculus*). The six most highly sampled taxonomic orders in the dataset (each with ≥ 44 species in the phylogeny; 571 species in total) were used to test for an effect of order on predicted mass regressed on observed mass. The model predicts adult mass with reasonable precision (*r*^2^ = 0.83) and accuracy (Figure 2B). Intercepts do not differ among orders and are not significantly different from 0 on a log^10^ scale (using Artiodactyla as a reference: intercept = −0.06, *t* = −0.3553, *P* = 0.7225). However, the slope depends on the taxonomic order (interaction effect: *F*_5,559_ = 2.8686, *P* = 0.0144), with slopes for the Artiodactyla, Carnivora, Chiroptera, and Rodentia being not significantly different from 1 (*P* > 0.40), and slopes for the Eulipotyphla and Primates being significantly less than 1 (*P* < 0.01). Therefore, the model provides accurate predictions for the Artiodactyla, Carnivora, Chiroptera, and Rodentia, but appears to be somewhat biased for the Eulipotyphla and Primates.

### Model predictions and macroevolutionary patterns

Although the model accurately predicts adult mass across more than seven orders of magnitude, it does not specify the sources of external selection on body size. These sources may be inferred from correlations of the tempo and mode of the evolution of body size with adaptive shifts or environmental, biogeographical, historical, and phylogenetic factors. To determine the effectiveness of life history theory in explaining macroevolutionary patterns, several well-defined patterns of mammal body size evolution associated with adaptive shifts, environmental changes, and biogeography were tested for effects on the adult mass predicted by the model. In these analyses, it is useful to consider the separate effects of the mean adult mortality rate (*μ*_*α*_ = 1/*E*_*α*_) and the reproduction rate in the model:

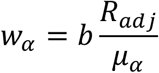

This version of the model shows that, all else being equal, optimal adult body mass will increase with a decreasing adult mortality rate or an increasing reproduction rate.

Mortality rates likely reflect levels of predation, and reproduction rates resource availability. The following analyses use data from the COMBINE database and the Life History dataset, unless stated otherwise.

### Foot structure and feeding mode

Changes in body size and other morphological traits of North American mammals during the Cenozoic Era (last 66 million years) have been linked to climate and associated vegetation changes ^38,59-62^. Climate cooling after the early Eocene epoch resulted in the spread of open woodland and savannah, followed by grasslands. It is proposed that in response, some herbivore lineages switched from browsing to grazing, with associated increases in unguligrade (hoofed) cursoriality and body size ^37^. The primary driver of increased size is argued to be cellulose fermentation of a low-quality diet by grazers, which requires a large body size for the rate of energy assimilation to exceed the rate of respiration^37,41^. Larger, faster grazers are argued to have selected for increases in digitigrade (toe-walking) cursoriality and size in carnivores, which allowed capture of a wide range of prey sizes ^37^. The larger, faster digitigrade carnivores are argued to have in turn constrained, through competition or predation, plantigrade (sole-walking) mammals from evolving large sizes ^37^.

The effects of feeding mode and cursoriality on the model’s predictions were tested using terrestrial sub-Saharan African mammals as an analog of North American mammal assemblages during the Miocene Epoch. Sub-Saharan Africa provides a reasonable analog because of its tropical rainforests and widespread tropical and subtropical open woodlands, savannahs, and grasslands, and because it did not experience megafaunal extinctions at the end of the Pleistocene ^40,51,60^. Density distributions confirm that although there is substantial overlap between categories, body mass tends to increase from plantigrade to digitigrade to unguligrade browser to unguligrade grazer (Figure 3A). However, there is no effect of foot structure/feeding mode on the prediction of adult mass by the model (Figure 3B). Nevertheless, the adult mortality rate is significantly lower for digitigrade species than other species of the same mass (intercept difference = 0.1378; *t* = 3.8432; *P* = 0.0004; Figure 3C). It is speculated that this effect is due to digitigrade species, which are mostly carnivorous, experiencing lower rates of predation. In contrast, there is no significant effect of foot structure/feeding mode on the relationship between reproduction rate and adult mass (Figure 3D). These patterns indicate that any effects of foot structure and feeding mode on the evolution of adult body mass occur through their effects on the adult mortality rate and the reproduction rate as specified by the model. For example, unguligrade species have evolved large sizes because of their low adult mortality rates and high reproduction rates. It may be speculated that these rates are due to high levels of cursoriality and resource availability, respectively.

**Figure 3.**
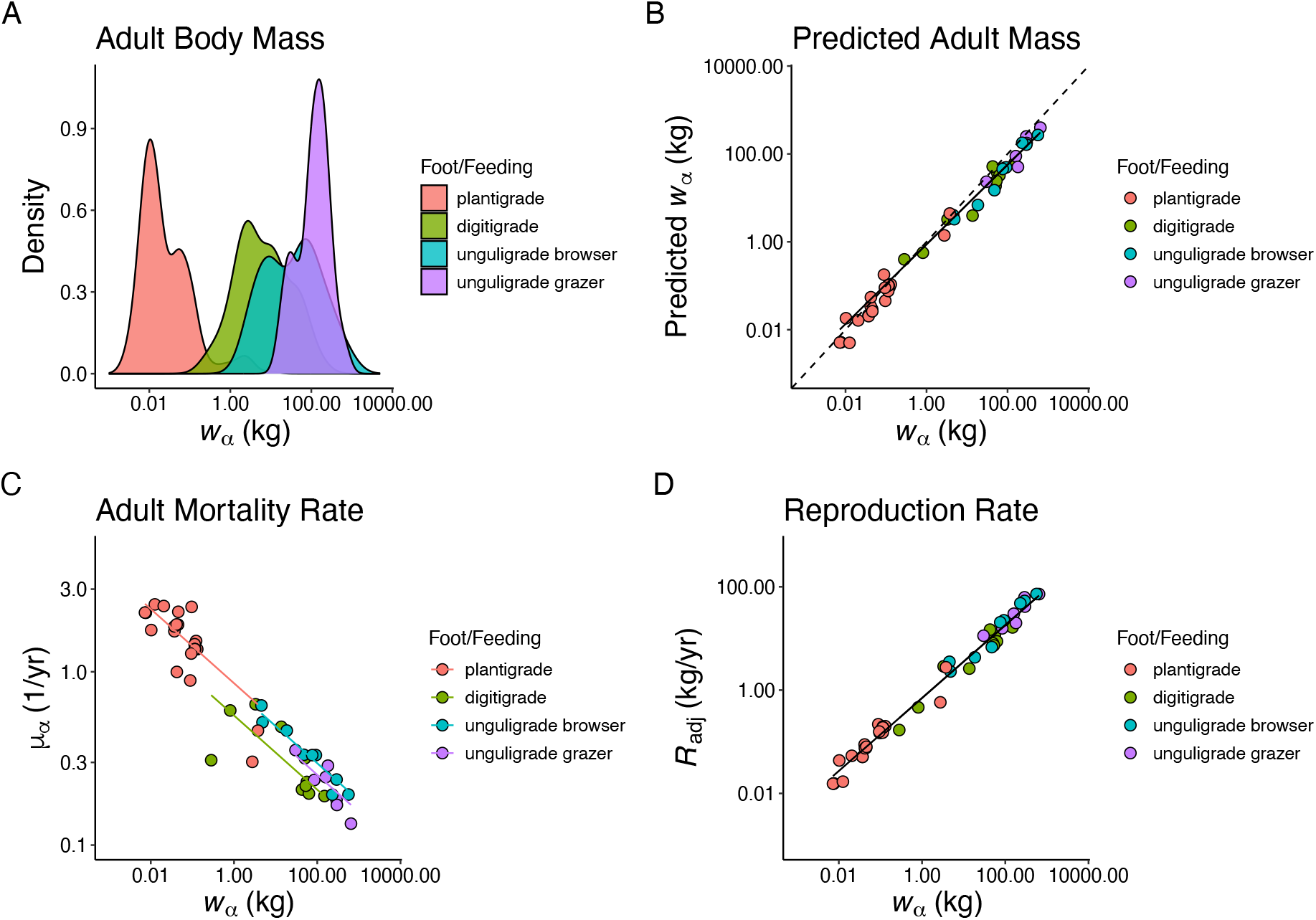
Terrestrial sub-Saharan African mammals: foot structure and feeding mode and the prediction of adult body mass. (A) Density distributions of adult body mass. (B) PGLS log-log regression of predicted adult mass on observed: λ = 1.00, κ = 0.89, δ = 3.00; r^2^ = 0.93, F_1,49_ = 676.5, P < 2.20 × 10^−16^; y = −0.06 + 0.91x. Dashed line indicates y = x. (C) PGLS ancova of foot structure and feeding mode on adult mortality rate with adult mass as a covariate: λ = 0.00, κ = 1.52, δ = 3.00; r^2^ = 0.95, F_4,46_ = 207.1, P < 2.20 × 10^−16^. (D) PGLS log-log regression of reproduction rate on adult mass: λ = 1.00, κ = 1.09, δ = 3.00; r^2^ = 0.91, F_1,49_ = 466.7, P < 2.20 × 10^−16^; y = −0.15 + 0.71x. Density plots are based on 363 species in seven orders. Predicted mass and life history plots are based on 51 species in seven orders.

### Aquatic living and feeding mode

Lineages transitioning to aquatic living show increases in body size ^42^, with maximum rates of evolution for whales (order Artiodactyla, infraorder Cetacea) being twice those of other lineages ^36^. Whales were compared to terrestrial artiodactyls to determine whether environment (aquatic or terrestrial) and feeding mode affect the prediction of adult body mass by the model. The semiaquatic hippopotamuses were excluded. Whales were categorized by their feeding mode as either toothed whales (parvorder Odontoceti) or baleen whales (parvorder Mysticeti). Terrestrial artiodactyls are primarily grazing or browsing herbivores. Whales are generally larger than terrestrial artiodactyls, and baleens whales are generally larger than toothed whales (Figure 4A). There is some effect of feeding mode on the prediction of adult mass by the model, with predicted mass of baleen whales higher than that of terrestrial artiodactyls and toothed whales after controlling for observed mass (main effect of environment/feeding mode: *F*_2,54_ = 5.2569, *P* = 0.0082; Figure 4B). Although, the model predicts mass very well overall (*r*^2^ = 0.90). There are also some effects of environment/feeding mode on the relationship between the adult mortality rate and adult mass, with mortality rates being generally lower for toothed whales compared to terrestrial artiodactyls and the sign of the relationship being possibly positive for baleen whales (interaction effect: *F*_2,52_ = 5.3409, *P* = 0.0078; Figure 4C). This may reflect lower rates of predation in aquatic environments. Toothed whales also have significantly lower, and baleen whales significantly higher, reproduction rates than terrestrial artiodactyls, controlling for observed body mass (main effect of environment/feeding mode: *F*_2,54_ = 11.951, *P* = 5.0440 × 10^−5^; Figure 4D). This may reflect differences in diet between toothed and baleen whales, with baleen whales utilizing highly productive food sources ^63,64^. The lower reproduction rate of toothed whales appears to cancel the effect of their lower mortality rate on predicted body mass since they do not differ from terrestrial artiodactyls in the relationship between predicted and observed mass. The higher reproduction rate of baleen whales may explain why their predicted mass is higher, closer to observed mass, than that of the other two groups. Therefore, aquatic living and diet appear to affect body size through their effects on adult mortality and the reproduction rate in ways consistent with the model.

**Figure 4.**
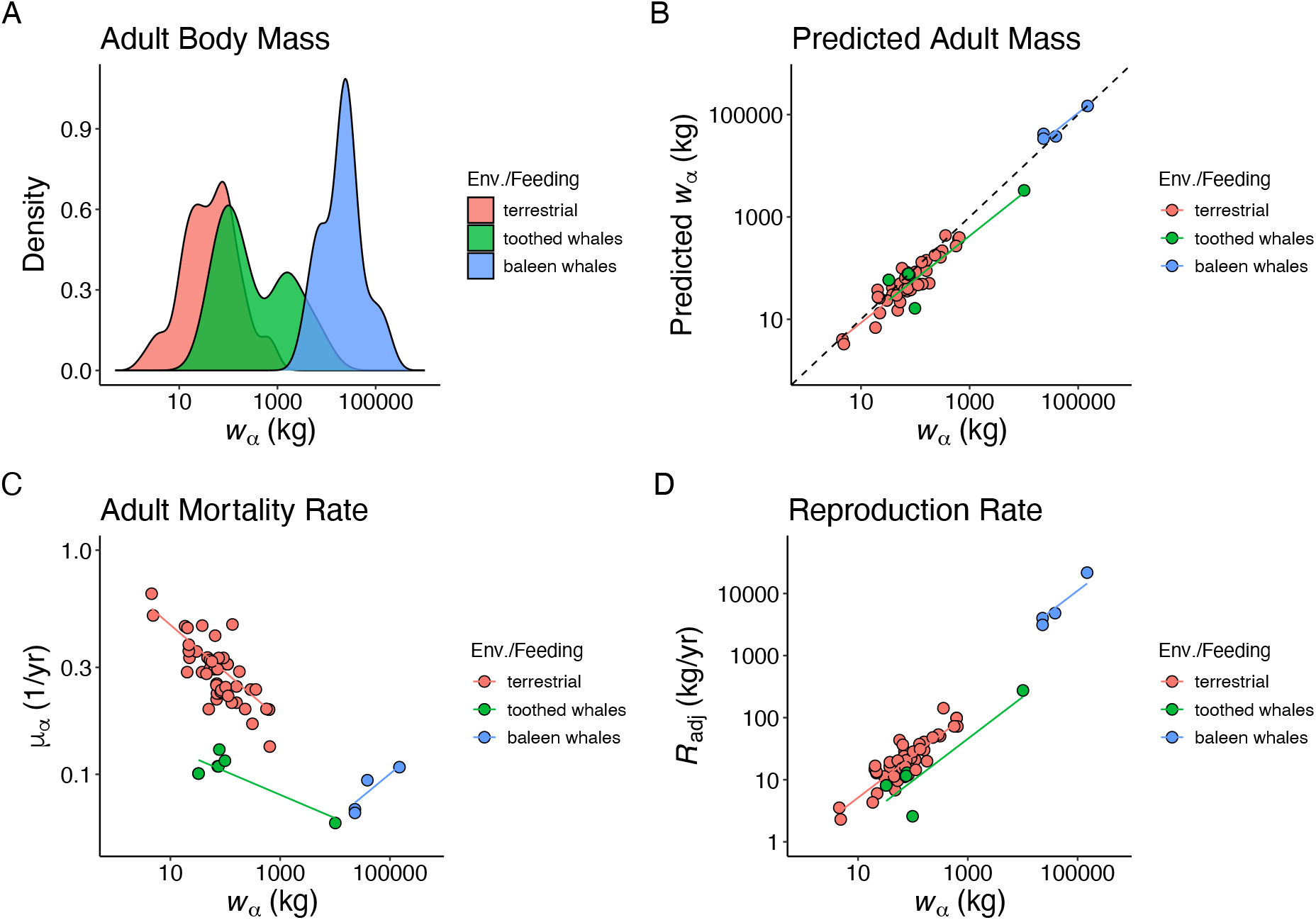
Whales and terrestrial artiodactyls: environment and feeding mode and the prediction of adult body mass. (A) Density distributions of adult body mass. (B) PGLS ancova of environment and feeding mode on predicted mass with observed mass as a covariate: λ = 0.95, κ = 0.88, δ = 3.00; r^2^ = 0.90, F_3,54_ = 167.6, P < 2.20 × 10^−16^. Dashed line indicates y = x. (C) PGLS ancova of environment/feeding mode on adult mortality with adult mass as a covariate: λ = 0.94, κ = 1.03, δ = 3.00; r^2^ = 0.73, F_5,52_ = 27.97, P = 1.237 × 10^−13^. (D) PGLS ancova of environment/feeding mode on the reproduction rate with adult mass as a covariate: λ = 0.94, κ = 1.02, δ = 3.00; r^2^ = 0.88, F_3,54_ = 133.8, P < 2.20 × 10^−16^. Density plots are based on 321 species. Predicted mass and life history plots are based on 58 species.

### Flight and diet

Bats (order Chiroptera), the only mammals with powered flight, are primarily insectivorous, with the main exception of fruit bats (family Pteropodidae), which feed primarily on nectar and fruit. The lineage leading to the common ancestor of bats exhibits an explosive rate of decrease in body size ^19^, whereas the lineage leading to fruit bats shows an accelerated increase in body size ^43^. To determine whether flight and diet affect predicted body mass, bats were compared to two orders, the primarily terrestrial and insectivorous Eulipotyphla (shrews, moles, hedgehogs, etc.) and the primarily arboreal and frugivorous or folivorous Primates (lemurs, tarsiers, monkeys, apes, etc.). The fossorial moles (family Talpidae) and armored hedgehogs (subfamily Erinaceinae) of the Eulipotyphla were excluded from analyses. This dataset consists of 2102 species with body mass data. There is almost complete overlap in the density distributions of mass between the Eulipotyphla species analyzed and the primarily insectivorous “microbats”, while fruit bats are generally larger, and primates generally larger still (Figure 5A). This pattern suggests that the overlap in mass between microbats and the Eulipotyphla is best explained if the rapid decrease in body size in the lineage leading to extant bats was in response to either their adoption of an insectivorous diet or to early adaptations to flight. The generally smaller size of fruit bats compared to primates may be due to metabolic and morphological constraints of flight.

**Figure 5.**
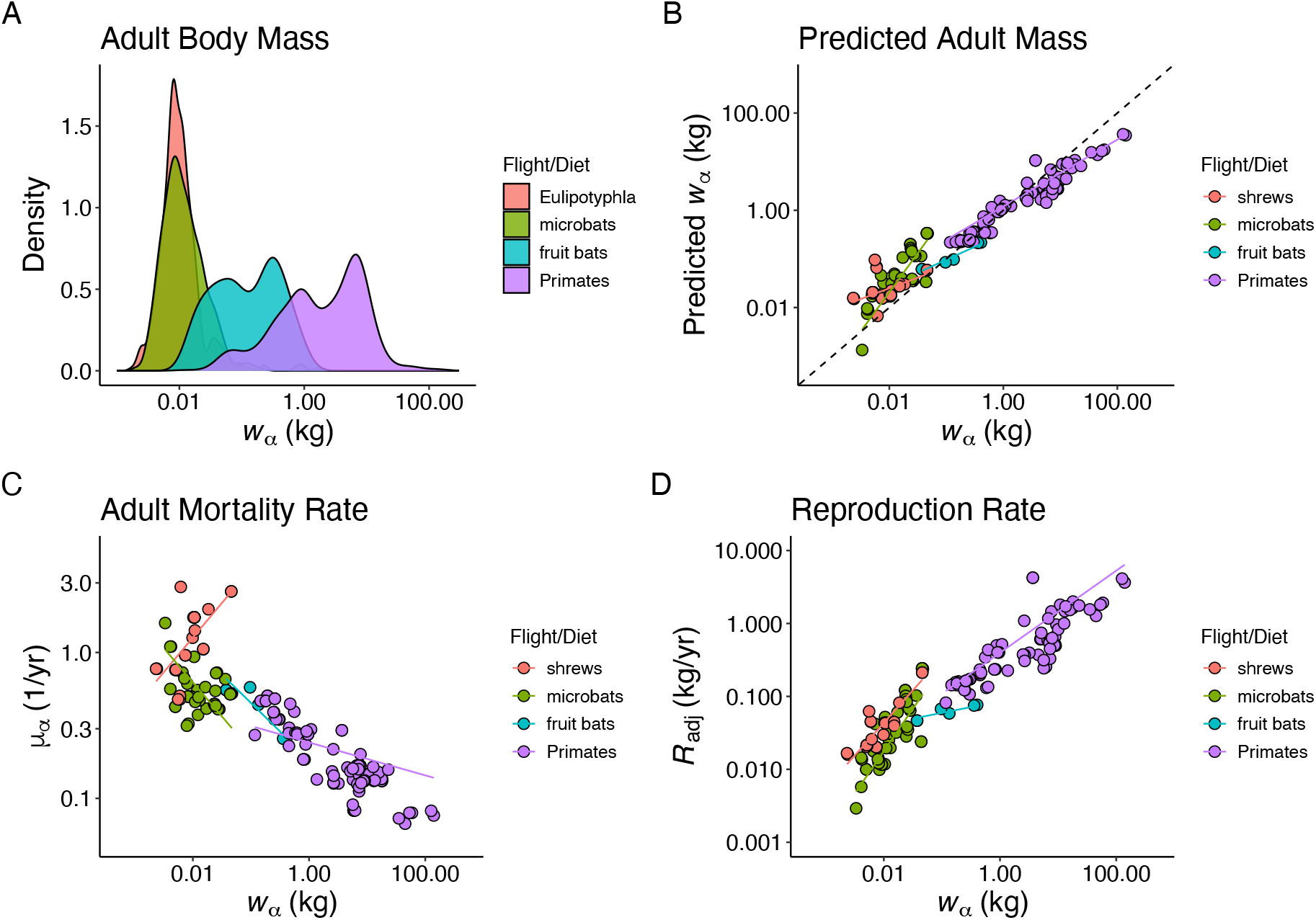
Bats, Eulipotyphla, and Primates: flight and feeding mode and the prediction of adult body mass. (A) Density distributions of adult body mass. (B) PGLS ancova of the effects of flight/diet on predicted adult mass with observed adult mass as a covariate: λ = 1.00, κ = 1.37, δ = 0.82; r^2^ = 0.66, F_7,176_ = 48.1, P < 2.20 × 10^−16^. Dashed line indicates y = x. (C) PGLS ancova of the effects of flight/diet on adult mortality rate with adult mass as a covariate: λ = 1.00, κ = 1.63, δ = 0.55; r^2^ = 0.36, F_7,176_ = 14.04, P = 2.042 × 10^−14^. (D) PGLS ancova of the effects of flight/diet on reproduction rate with adult mass as a covariate: λ = 1.00, κ = 1.42, δ = 0.91; r^2^ = 0.61, F_7,176_ = 38.9, P < 2.20 × 10^−16^. Density plots are based on 2102 species. Predicted mass and life history plots are based on 184 species.

Data on rates of adult mortality and reproduction, allowing body mass to be predicted by the model, are available for only 184 of the species in this dataset, of which the Eulipotyphla are represented exclusively by shrews (family Soricidae). There is an effect of flight/diet on the prediction of adult mass by the model: microbats have a slope of predicted mass regressed on observed mass significantly greater than one (1.61; *t* = 3.03901, *P* = 0.0014), whereas slopes for the other taxa are less than one (shrews: 0.44; *t* = −3.7206, *P* = 0.0003) or not significantly different from one (fruit bats and primates: *P* > 0.8) (interaction effect: *F*_3,176_ = 15.8644, *P* = 3.565 × 10^−9^; Figure 5B). In the case of the adult mortality rate regressed on adult mass, shrews are unusual in exhibiting a positive relationship (interaction effect: *F*_3,176_ = 26.1452, *P* = 4.905 × 10^−14^; Figure 5C). Reproduction rate is positively related to adult mass for all taxa, but slopes differ (interaction effect: *F*_3,176_ = 8.6715, *P* = 2.131 × 10^−5^; Figure 5D). Microbats appear more like shrews than fruit bats in terms of reproduction rate, both exhibiting high positive slopes. The model tends to overestimate mass for shrews and microbats. This may be due to the use of maximum growth rate as an analog of reproduction rate in the model since both shrews and small bats in temperate environments reach maximum mass before reaching sexual maturity ^65,66^ and thus may have exceptionally high growth rates. Otherwise, the model performs reasonably well (*r*^2^ = 0.66). Overall, flight and diet appear to affect predicted mass through their effects on adult mortality and reproduction rate. For example, shrews and microbats are small because they have high mortality rates and low reproduction rates, primates are larger because of their lower mortality rates and higher reproduction rates, and fruit bats are of intermediate size, reflecting their intermediate rates.

### Island endemicity

To test the effects of island endemicity on the prediction of adult body mass by the model, terrestrial (non-volant and non-marine) species on isolated islands were matched to mainland sister species. The island rule is upheld. Using data from the PHYLACINE database ^32^, regression of the adult mass of island species (*w*_*αI*_) on their matched mainland species (*w*_*αM*_) gives a slope (0.79) that is significantly less than 1 (*t* = −3.4100, *df* = 54, *P* = 6.1805 × 10^−4^; Figure 6A), indicating that small mainland species tend to evolve larger size on islands and large mainland species tend to evolve smaller size on islands. Nevertheless, the model predicts accurately the adult mass for both island and mainland species for which data in the Life History dataset are available (Figure 6B). The main causal explanation for insular gigantism is ecological release from predators or competitors ^47^. The model predicts that this will be caused by either a decrease in the adult mortality rate, due to the absence of predators, or an increase in the reproduction rate, due to the absence of competitors, or both. This prediction was tested by calculating log ratios of island (*I*) to matched mainland (*M*) species for adult mortality rate 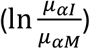 and reproduction rate 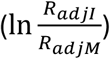 and comparing these to the log ratio of their adult masses 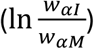 for the six species pairs for which this was possible. A positive log ratio indicates a higher value for the island species, whereas a negative log ratio indicates a lower value for the island species. The model prediction is upheld in that giant island species (positive log mass ratios) have negative log mortality rate ratios (Figure 6C) and positive log reproduction rate ratios (Figure 6D). Therefore, for the two species indicated, insular gigantism is consistent with both a lower adult mortality rate and a higher reproduction rate, as predicted by the model.

**Figure 6.**
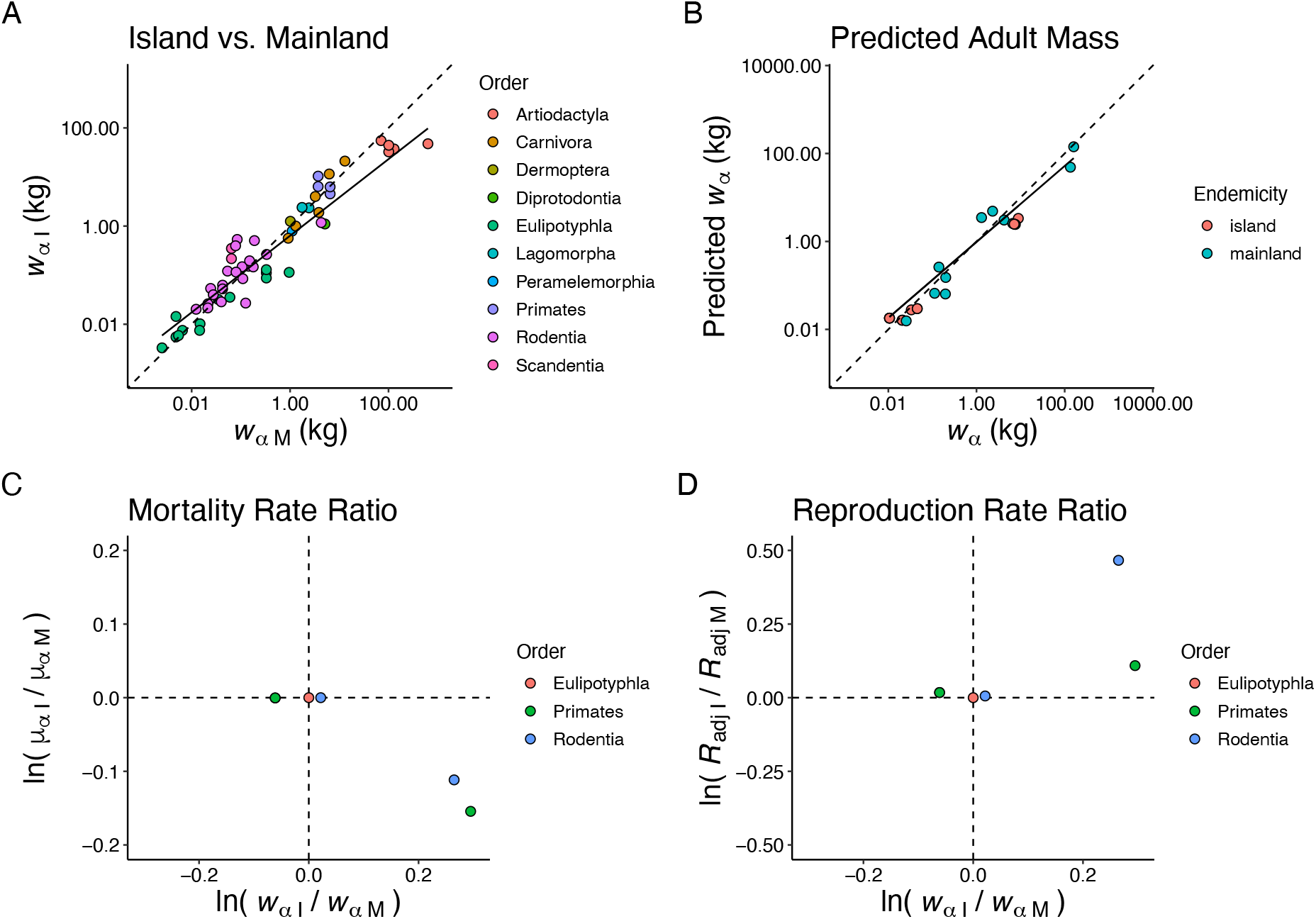
Island endemicity and the prediction of adult body mass. (A) Test of the island rule. PGLS log-log regression of island (w_αI_) on mainland (w_αM_) mass for 56 species pairs in 10 orders: λ = 0.94, κ = 1.00, δ = 3.00; r^2^ = 0.76, F_1,54_ = 172.8, P < 2.20 × 10^−16^; y = −0.16 + 0.79x. (B) PGLS log-log regression of predicted adult mass on observed for island and mainland species: λ = 0.67, κ = 3.00, δ = 0.20; r^2^ = 0.90, F_1,23_ = 200.5, P = 7.61 × 10^−13^; y = −0.01 + 0.86x. Dashed line indicates y = x. (C) Test of the life history model: log mortality rate ratio plotted against log mass ratio. (D) Test of the life history model: log reproduction rate ratio plotted against log mass ratio.

## Discussion

The model’s success is remarkable considering the variation in adult mass, physiology, life history, and ecology represented by the 22 orders analyzed. This success may be attributed in part to all mammals, and perhaps all animals, having fundamentally the same metabolic constraints on productivity ^6,53,67^. It appears that the diversity of body size among mammals can be explained by life history optimization with respect to adult mortality under metabolic constraints on productivity. That is, mammals grow to a size at which their reproduction rate over their reproductive lifespan maximizes fitness.

Nevertheless, the tests of the model are correlational, which leaves open the question of causality. For example, does an increase in the adult mortality rate, all else being equal, select for a decrease in size, as predicted by the model, or does a decrease in size increase the adult mortality rate? However, a classic experiment with the fruit fly *Drosophila* ^68^, which also exhibits deterministic growth, confirms this key prediction of the model. In this experiment, populations were maintained at constant density (constant resource availability) and those subject to high adult mortality rates evolved decreases in lifespan and ages and sizes at maturity compared to populations subject to low mortality rates.

As the model stands, the true reproduction rate is estimated as 0.35 of the maximum production rate, estimated as the maximum growth rate. Ideally, the reproduction rate would be estimated directly from the total cost of reproduction, rather than from offspring mass at independence, as is currently possible, because indirect costs of reproduction (other than energy in offspring) account for the majority of reproduction costs in mammals ^57^. In addition, indirect costs appear to account for a higher proportion of the total cost of reproduction in larger species ^57^. Furthermore, the cost of somatic maintenance will constrain the proportion of total production allocated to reproduction, and this cost may also increase with body size. Both susceptibility to infection or parasitism and cancer mortality are expected to increase with body size, but both bacteria killing capacity ^69^ and cancer mortality rate ^70^ do not vary with size in mammals but see ^71^. This suggests that larger species invest disproportionately more in antibacterial defences and cancer suppression, and perhaps in somatic maintenance in general ^58^. Extending the model to incorporate the true costs of reproduction and somatic maintenance should provide more accurate predictions. However, whether such models can be tested will depend on the availability of appropriate data.

Coulson, et al. ^1^ have used a different life history modelling approach to predict optimal size in species with determinate growth and density-dependent population regulation. They use a set of recursion equations relating body size to survival, reproduction, growth, and offspring size, each a function of population density, and iterate these to generate population dynamics, with density dependence acting either through juvenile survival or fecundity. Trade-offs between life history traits are generated by population regulation. They repeat this procedure for different life history strategies, each represented by a growth trajectory and a size at sexual maturity, and compare the fitnesses of the strategies. The model has not been tested because of the numerous parameters that must be estimated. However, the model’s main prediction, that selection on body size is disruptive, resulting in either gigantism or dwarfism, is untenable (this is unrelated to the island rule, which predicts that body masses converge toward an intermediate value on islands). The model’s behavior is attributed to the use of population carrying capacity as a measure of fitness ^1^.

Carrying capacity has been argued to be the appropriate measure of fitness when *selection* is density dependent (population density imposes selection directly or indirectly) and demography changes stochastically ^72^. The evidence for density dependent selection is mostly in the form of phenotypically plastic correlates of population density, which are presumed to be adaptive ^73-76^. Nevertheless, a simple thought experiment shows that carrying capacity cannot be a general measure of fitness in persistent populations. If predators select for better camouflage in a prey species, but population density of the prey species is limited by food supply and camouflage is uncorrelated with food acquisition or assimilation, then better camouflage will evolve regardless of density and have no effect on equilibrium density. That is, genotypes that encode better camouflage do not in any sense have higher carrying capacity. The insufficiency of carrying capacity as a measure of fitness is also demonstrated in the classic experiment with fruit flies described above. Even though larvae and adults were maintained at constant population densities, populations subject to high adult mortality evolved decreases in lifespan and ages and sizes at maturity compared to populations subject to low mortality ^68^.

The model presented here assumes that population density is regulated through juvenile survival (Supplementary Information) and therefore fitness is measured as lifetime reproductive success. With density-independent selection in a population, evolutionary invasion (evolutionary stable strategy, ESS) analyses show that the appropriate measure of fitness depends on the life stage that is the target of density dependence ^77^. With population size regulated through fertility, juvenile survival, adult survival, or the age of maturity, fitness is proportional to lifetime reproductive success.

The model helps explain how extrinsic factors (e.g., climate and associated vegetation changes, diet, feeding mode, cursoriality, aquatic living, flight, and island endemicity) may affect the macroevolution of body size. The effects of extrinsic factors may be understood through their effects on the rates of adult mortality and reproduction in the model. Generally, small species have high mortality rates and low reproduction rates, whereas large species have low mortality rates and high reproduction rates. For example, the largest unguligrade mammals may have low adult mortality rates because of their cursoriality and high reproduction rates because they utilize seemingly unlimited resources. Similarly, baleen whales appear to have evolved large sizes because of low adult mortality rates associated with aquatic living and high reproductive rates associated with highly productive food sources. And microbats may have evolved sizes similar to those of shrews because of similar high mortality rates and low reproduction rates, possibly associated with a primarily insectivorous diet or other constraints on body size affecting mortality and reproduction rates.

This study contributes to our understanding of the links between micro- and macroevolution ^25^. Macroevolutionary-scale phenomena such as species selection associated with the end Cretaceous mass extinction or Pleistocene megafauna extinctions clearly affect distributions of body size ^78^. However, in the background are microevolutionary process captured by the model presented here. These results suggest that external factors affecting both adult mortality and productivity determine the constraints to which life histories are optimized. Thus, regardless of whether the rate of body size evolution changes over time or among lineages, or whether body size evolves in response to new niches, environmental change, or biogeographical factors, body size is mostly evolving incrementally and continuously towards changing optima as dictated by the principles of life history theory.

## Methods

### Taxonomy

Taxonomy was harmonized between datasets following the guidelines of Grenié, et al. ^79^. The Global Biodiversity Information Facility (GBIF) ^80^ was used to resolve species names in datasets using the R package *rgbif* ^81,82^. The Mammal Diversity Database version 1.10 ^83^, which contains taxonomic information for all 6,615 known extant (and recently extinct) mammal species at the time of writing ^84^, was used as the taxonomic reference. In harmonizing the taxonomy of datasets, species names may have changed, and species numbers may have been reduced due to species synonyms.

### “Demography dataset”

The “Demography dataset” contains variables whose values were computed from demographic data from natural, undisturbed populations of mammals. The main variables are the age of female first reproduction (*α*), probability of survival from birth to maturity (*S*_*α*_), life expectancy at maturity (*E*_*α*_), and annual fecundity in female offspring (*m*). Forty-three species of mammal with all of these data are a subset of 64 species from Purvis and Harvey ^85^ after omitting domesticated species and species with missing values. Values were also estimated from matrix population models (MPMs) in the COMADRE database version 4.23.3.1 ^86^ using the *Rcompadre* and *Rage* R packages ^87^. MPMs were filtered for basic validity criteria ^87^ and then filtered further to include only those matching all of the following criteria: (1) pooled studies of females in the wild, (2) unmanipulated populations, (3) age classes at intervals of one year, and (4) divided survival and fecundity matrices. MPMs for only 10 species matched these stringent criteria. Values for species in common between the two datasets were averaged, giving a total of 49 species from eight orders of mammal. Litter size, age at weaning and adult, neonate, and weaning mass were added from the COMBINE database of mammal life history and ecological traits ^30^ (see below). With these added variables there are 41 species in eight orders with complete data.

### “Life History dataset”

Life history data are from the COMBINE database ^30^, a curated coalescence of several databases (e.g., ^32,34^) containing 5,808 species after taxonomic harmonization (88% of recognized species). This database pools data on mammal life history and extrinsic (ecological and biogeographical) traits from all sources published between 1999 and 2020. Variables from the database used here are age at weaning, female age at maturity, gestation length, litter size, number of litters per year, and adult, neonate, and weaning mass. Age of female first reproduction was calculated as the sum of female age at maturity and gestation length as this was found to give more reliable results (fewer outliers) than the reported age at first reproduction. Values for all of these variables, without missing data, are available for 794 species in 22 of the 27 orders of extant mammals ^84^.

### Phylogenetic comparative analyses

Statistical relationships among life history variables across species corrected for phylogenetic non-independence were estimated using phylogenetic generalized least squares (PGLS) analyses ^88^ as implemented by the R package *caper* ^89^. This method estimates the covariance between each pair of taxa using the branch lengths of a time-scaled (ultrametric) phylogeny. The initial covariance matrix is estimated assuming a Brownian motion model of evolution, in which phenotypes track adaptive optima that change incrementally and continuously in an unbiased random walk. To account for departures from strict Brownian motion, transformations are applied to improve the fit of the model to the data using maximum likelihood methods. Internal branch lengths are multiplied by *λ*, such that *λ* = 0 indicates the evolution of traits independent of the phylogeny, whereas *λ* = 1 indicates evolution fully consistent with Brownian motion on the phylogeny. Values of the covariance matrix are also raised to the power *δ*, which is a transformation of the sum of the lengths of the shared branches between two taxa, such that *δ* = 1 indicates gradual evolution consistent with Brownian motion, whereas *δ* < 1 indicates initial rapid evolution as might occur with an adaptive radiation. And all branch lengths are raised to the power *κ*, such that *κ* = 1 indicates anagenesis (change along branches) consistent with Brownian motion, whereas *κ* → 0 indicates cladogenesis (change occurs when branches split). Values for these parameters are reported with the results of individual analyses. The evolution of mammal body size is better described by biased Brownian motion than by alternatives such as “pulsed change” models or “adaptive landscape” models, including the Ornstein-Uhlenbeck model ^20^.

For this purpose, the most up-to-date fully resolved and accurately time-scaled molecular phylogeny available for mammals ^31^, containing 3,896 species after taxonomic harmonization (59% of recognized species), was used. The methods used to infer this phylogeny overcome previous limitations that lead to imprecise and inaccurate node ages estimated with “backbone-and-patch” methods ^90^. The phylogeny includes all 41 species in the Demography dataset and 714 species in 21 orders in the Life History dataset.

### Testing the life history optimisation model

### Production rate scaling exponent

The production rate scaling exponent was set to *b* = 0.73 based on the allometry of basal metabolic rate (BMR) across 549 species of mammal for which reliable BMR values are available ^91^. This value matches closely the mean *intraspecific* scaling exponent for BMR measured over large mass ranges across animals (*b* = 0.74; 95% profile likelihood confidence interval for the mean: 0.699, 0.777) ^6^.

#### Adult mortality rate

Life expectancy at maturity, *E*_*α*_ (and therefore the mean adult mortality rate, *μ*_*α*_ = 1/*E*_*α*_), which is difficult to measure in the wild and thus rarely reported, is not reported in the COMBINE database. Therefore, for the Life History dataset, *E*_*α*_ was estimated from its well-established strong log-log linear relationship with the age at first reproduction, *α*, ^92^ using the Demography dataset (PGLS regression: *λ* = 0.81, *κ* = 1.07, *δ* = 3.00; *r*^2^ = 0.81, *F*_1,47_ = 201.6, *P* < 2.2 × 10^−16^; *y* = 0.25 + 0.90*x*; Supplementary Information). As *α* is not contained in the model, it provides an independent estimate of *E*_*α*_. Formal imputation methods are not applicable to estimating *E*_*α*_ since it is available for only 6% of species in the Life History dataset and imputation methods require at least 40% non-missing data for reliable results ^93^.

#### Growth rate

Individual maximum growth rate was estimated from the *w*_0_-parameterization of the Gompertz growth equation ^94^:

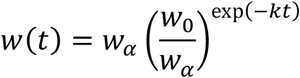

where *w*(*t*) is mass at age *t, w*_0_ is neonate mass, and *k* is the growth coefficient. This model generally provides a better fit to the growth of mammals than either the von Bertalanffy or the logistic growth models and its parameters have clearer biological meanings ^95^. *k* was estimated by setting *w*(*t*) and *t* to the mass and age at weaning, *w*_w_ and *t*_w_, and solving for *k*:

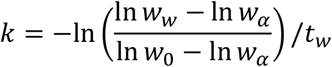

The maximum growth rate is calculated as ^94^: *k*_max_ = *kw*_*α*_/*e*.

### Foot structure and feeding mode

Data on foot structure and feeding mode for mammal species endemic to sub-Saharan African are from Skinner and Chimimba ^96^ and Kingdon, et al. ^97^. Species found only on Madagascar, or any other island, were excluded to avoid island effects on body size ^47^. As all unguligrade mammals are terrestrial, only terrestrial mammals were included to avoid effects of other modes of living on body size. Aquatic, arboreal, scansorial, flying, gliding, and fossorial species were excluded. Armoured species (hedgehogs, pangolins and porcupines) were also excluded due to potential effects of anti-predator adaptations on body size. Species with a saltatory mode of locomotion (e.g., rabbits, hares, springhares, elephant shrews, and gerbils) were excluded because their foot posture changes with mode of locomotion. All African members of the orders Artiodactyla and Perissodactyla are unguligrade except for hippopotamuses and rhinoceroses. For the order Carnivora, canids, felids, and hyaenids are digitigrade, as are some mustelids and herpestids. The aardvark is digitigrade (order Tubulidentata). Other mustelids and herpestids are plantigrade, as are all hyraxes (order Hyracoidea), most rodents (order Rodentia), and all shrews and their relatives (order Eulipotyphla). Most unguligrade species are herbivorous, primarily browsers or grazers. Most digitigrade species are carnivorous, while plantigrade species may exhibit various forms of herbivory, insectivory, or carnivory. These data were collected for 363 species in the seven orders mentioned above and for which body mass was available. Data from the Life History dataset, used to predict optimal body mass, are available for 54 of these species in seven orders in the phylogeny.

### Island endemicity

Data used to test the effects of island endemicity on body size are from the PHYLACINE database (Version 1.2.1) ^32^. Data from a large metanalysis test of the island rule ^47^ were not used because these are primarily from matched island-mainland conspecific populations and the necessary life history data are not available for the specific island populations. The PHYLACINE database contains data on body mass, island endemicity, and other traits for 5425 species of mammal after taxonomic harmonization. Species were classified as island endemic if they occurred *only* on isolated islands, separated from the mainland by water deeper than 110 m (not connected to the mainland during the last glacial maximum). Special care was taken when harmonizing the database taxonomy because 20 populations of terrestrial mammals on isolated islands identified as distinct species in the database are now considered as synonymous with species on land bridge islands (separated from the mainland by water ≤ 110 m deep) or the mainland ^84^. In these cases, the isolated island entry was deleted because data in the Life History dataset correspond to the more common and widespread population. Island species were matched with their potential mainland sister species by identifying their sister taxon in the phylogeny ^31^. If the sister taxon was a clade, then one species in the clade for which data are available in the Life History dataset was chosen at random, if available, as the match. Species on very large islands (e.g., Madagascar) were excluded by the fact that their sister taxon was invariably another isolated island species. Seals (Order Carnivora, Superfamily Phocoidea) and bats (Order Chiroptera) were excluded from analyses because of their enhanced mobility between islands and the mainland. This produced a dataset of 56 island-mainland species pairs in 10 orders. Of these, data from the Life History dataset are available for both members of only six species pairs in three orders.

