## Supplementary Information for "Life History Optimization and the Macroevolution of Mammal Body Size"

### Adult mortality rate

For the Life History dataset, $E_\alpha$ was estimated from its well-established strong log-log linear relationship with the age at first reproduction, $\alpha$, [1] using the Demography dataset (PGLS regression: $\lambda = 0.81, \kappa = 1.07, \delta = 3.00;\ r^2 = 0.81, F_{1,47} = 201.6, P < 2.2 \times 10^{-16}; y = 0.25 + 0.90x$; Figure 1).

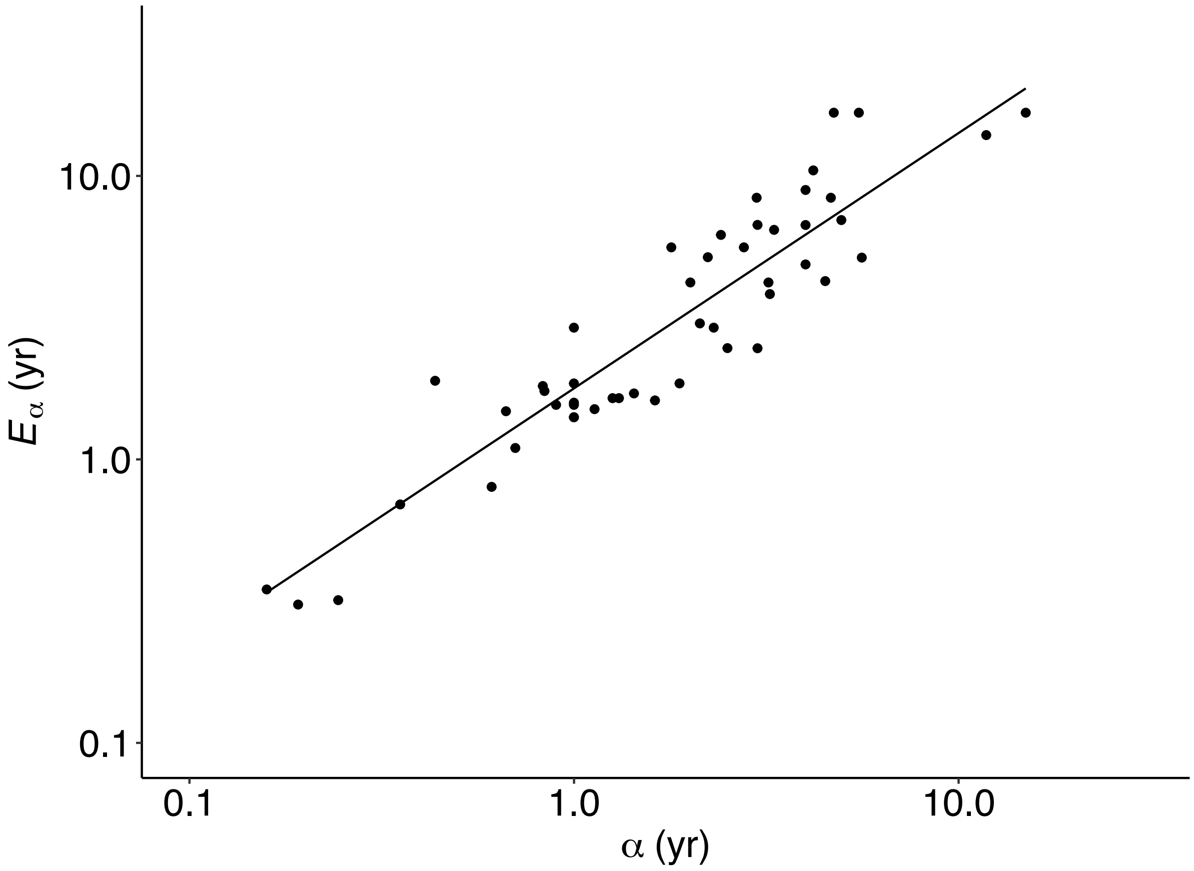


*Figure 1. PGLS log-log regression of life expectancy at maturity, $E_\alpha$, on age at first reproduction, $\alpha$, for 49 species in 8 orders ($\lambda = 0.81, \kappa = 1.07, \delta = 3.00;\ r^2 = 0.81, F_{1,47} = 201.6, P < 2.2 \times 10^{-16}; y = 0.25 + 0.90x$).*

### Testing the model's assumptions

#### Population stability

The model's most basic assumptions are that the population is stable and that density-dependent regulation occurs through juvenile recruitment to the adult age class [2]. Density-dependent population regulation is ubiquitous in mammals and other animals [3-5] and generally acts through changes in fecundity or juvenile survival [6-11]. These assumptions were tested using the following framework. The net reproductive rate of a population, the mean lifetime reproductive success of females, is equal to one for a stable population:

$$R_0 = \sum_{x=\alpha}^{\omega} l_x \, m_x = 1$$

where $l_x$ is the probability of surviving from birth to age $x$, and $m_x$ is fecundity (number of female offspring) at age $x$, $\alpha$ is the age of first reproduction, and $\omega$ is the age of last reproduction [12]. This may be rewritten in terms of the demographic variables that are typically studied:

$$R_0 = S_\alpha E_\alpha m_f = 1$$

where $S_\alpha$ is the probability of surviving from birth to sexual maturity (juvenile survival), $E_\alpha$ is life expectancy at maturity, and $m_f$ is the mean rate of fecundity in female offspring [2,13-15]. Therefore, in a stable population, the rate of juvenile recruitment to the adult age class, $S_\alpha m_f$, is expected to equal the mean adult mortality rate, $\mu_\alpha = 1/E_\alpha$.

Population stability is confirmed for demographic data from 49 species of mammal in eight orders: the rate of juvenile recruitment to the adult class, $S_\alpha m_f$, is approximately equal to the mean adult mortality rate, $1/E_\alpha$, across species (Figure 2A). Regression of $S_\alpha m_f$ on $1/E_\alpha$ shows that the intercept of the regression line is not significantly different from 0 on a log-log scale (intercept = 0.05; $t = 0.928,\ df = 47,\ P = 0.36$) and that the slope is not significantly different from 1 (slope = 1.07; $t = 0.875, df = 47, P = 0.39$).

The target of population regulation may be deduced from the relationships between the components of the net reproductive rate. Although there is a strong relationship between the rate of fecundity ($m_f$) and the adult mortality rate across species (Figure 2B), this is unlikely to explain population stability *within* species since such a relationship across species is indicative of adaptation, presumably due to life history optimization. Fecundity and life expectancy are evolved traits whose values will vary around species means. For example, placing species in zoos will not change this relationship [16-18]. The probability of juvenile survival to maturity ($S_\alpha$), on the other hand, is a demographic variable whose value may vary from zero to one within a population for any species depending on environmental conditions. That is juvenile survival varies on a much shorter timescale, within a generation, than life history adaptation, which occurs over generations. The lack of a relationship between juvenile survival and adult mortality *across* species (Figure 2C) is consistent with juvenile survival being a target of population regulation *within* species. Juvenile survival also has no significant relationship with fecundity across species (Figure 2D). These results suggest that population regulation in mammals typically occurs through changes in juvenile survival rather than fecundity [2]. Values of fecundity are transformed by juvenile survival to give juvenile recruitment that is approximately equal to adult mortality for each species.

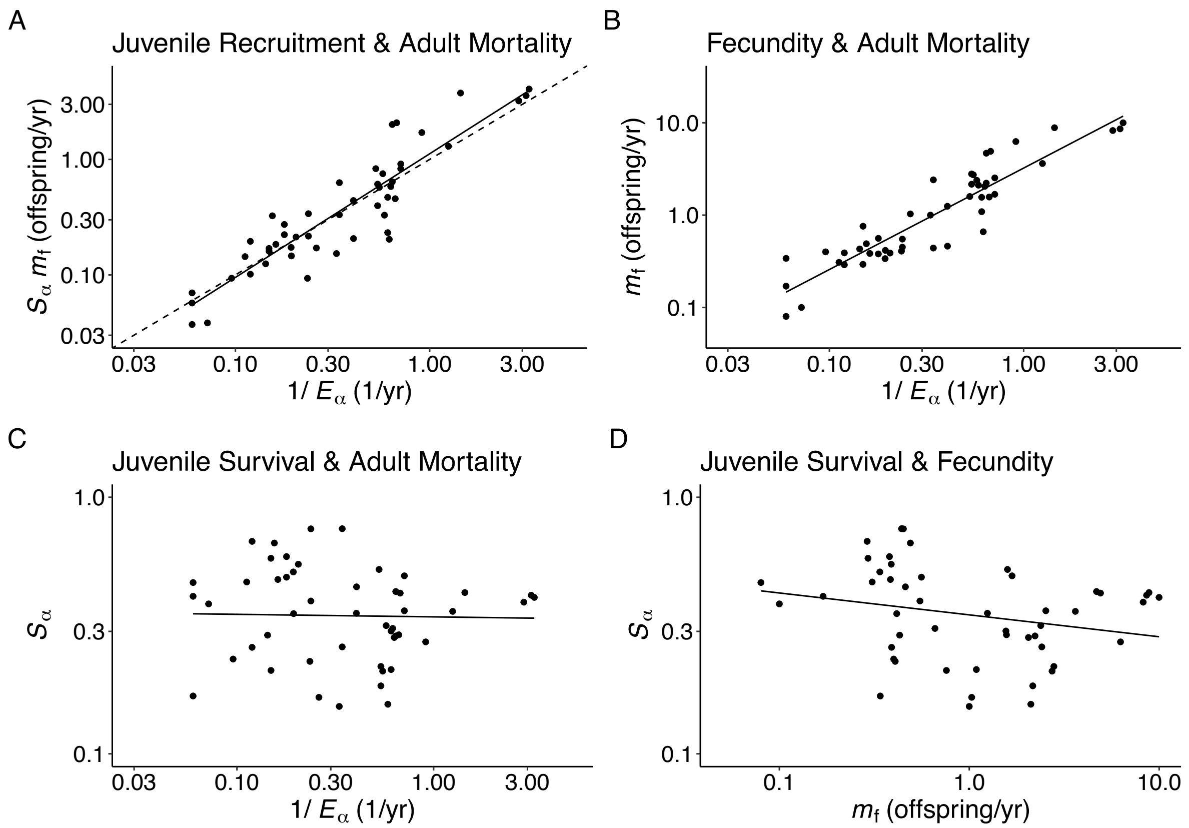


*Figure 2. PGLS log-log regressions of recruitment rate and its components for 49 species in 8 orders. (A) Recruitment rate, $S_\alpha m_f$, on adult mortality rate, $1/E_\alpha$ : $\lambda = 0.76, \kappa = 1.03, \delta = 3.00; r^2 = 0.79, F_{1,47} = 175,\ P < 2.20 \times 10^{-16}; y = 0.05 + 1.07x$. Dashed line indicates $y = x$. (B) Rate of fecundity (female offspring), $m_f$, on adult mortality rate: $\lambda = 0, \kappa = 1.08, \delta = 3.00; r^2 = 0.84, F_{1,47} = 248.7,\ P < 2.20 \times 10^{-16}; y = 0.51 + 1.10x$. (C) Probability of survival to maturity, $S_\alpha$, on adult mortality rate: $\lambda = 0.83, \kappa = 1.04, \delta = 2.20; r^2 = 0.00041, F_{1,47} = 0.01935,\ P = 0.89; y = -0.47 - 0.0099x$. (D) Probabiity of survival to maturity on rate of fecundity (female offspring): $\lambda = 0.83, \kappa = 1.01, \delta = 2.12; r^2 = 0.046, F_{1,47} = 2.279,\ P = 0.14; y = -0.46 - 0.09x$.*

### Fitness is lifetime reproductive success

Density-dependent regulation acting through juvenile survival supports another key assumption: that fitness is measured as lifetime reproductive success. With density-independent selection in a population, evolutionary invasion (evolutionary stable strategy, ESS) analyses show that the appropriate measure of fitness depends on the life stage that is the target of density dependence [19]. With population size regulated through fertility, juvenile survival, adult survival, or the age of maturity, fitness is proportional to lifetime reproductive success.

### Body size is optimized to adult mortality

A simplifying assumption, made for analytical tractability, is that mortality rate is independent of size around the time of maturity within species [20-22]. In the model, the adult mortality rate is an independent variable to which body size is optimized within the metabolic constraints of the production rate. Although mortality rate appears to decrease with increasing juvenile size in some populations of some species [23], limited evidence suggests that mortality is independent of size near the age of maturity [22]. In addition, the probability of survival to adulthood, $S_\alpha$, is independent of neonate, weaning, or adult mass, and of adult mortality, with a mean of 0.37 across 49 species in 8 orders in the Demography dataset. Finally, making mortality rate strictly size dependent has little effect on the optimal size or age at maturity with realistic parameter values [24]. More importantly, if population regulation acts through juvenile survival, then even if juvenile mortality depends on body size at a given population density, juvenile mortality will still be determined mainly by population density and thus will vary on a much shorter timescale (within a generation) than life history adaptation (across generations).

### Adjusting the reproduction rate

Reproduction rate was calculated as the product of the total fecundity rate (number of all offspring per unit time) and offspring mass at independence: $R = mw_i$. Following Charnov, et al. [25], offspring mass at independence is weaning mass, $w_w$, if offspring survive to weaning, otherwise it is their mass at death. Assuming that offspring that die before weaning have a mass that is the average of neonate and weaning masses, offspring mass at independence is these masses weighted by preweaning survival, $S_w$ [25]:

$$w_i = S_w w_w + (1 - S_w)(w_0 + w_w)/2$$

Since preweaning survival is available for few species, Charnov, et al. [25] estimate preweaning survival from it regression on litter size, $L$, for 13 species (5 artiodactyls and 8 rodents), giving the equation $S_w = 0.7L^{-0.35}$. This equation was used in the estimation of $w_i$. $S_w$ is expected to decrease with litter size in stable populations if population density is regulated by juvenile survival.

The reproduction rate, based on the mass at offspring independence, is expected to underestimate the true reproduction rate because indirect costs (energetic costs other than the energy content of offspring) account for the majority of reproduction costs in mammals [26]. As expected, using $R$ in the model ($w_\alpha = bE_\alpha R$) generally underestimates adult body mass, especially for large species (Figure 3A). However, the true value of $R$ is expected to be lower than the maximum production rate, $P_{max}$, because adults must invest in somatic maintenance. $P_{max}$ may be estimated as the maximum growth rate, $k_{max}$, from the Gompertz growth equation (see Methods). As expected, using $k_{max}$ in place of $R$ in the model overestimates adult body mass (Figure 3B). Therefore, the true reproduction rate is assumed to be some proportion of $k_{max}$. Using an adjusted reproduction rate of $R_{adj} = 0.35k_{max}$ in the model ($w_\alpha = bE_\alpha R_{adj}$) gives a good match between predicted and observed adult body mass (Figure 3C).

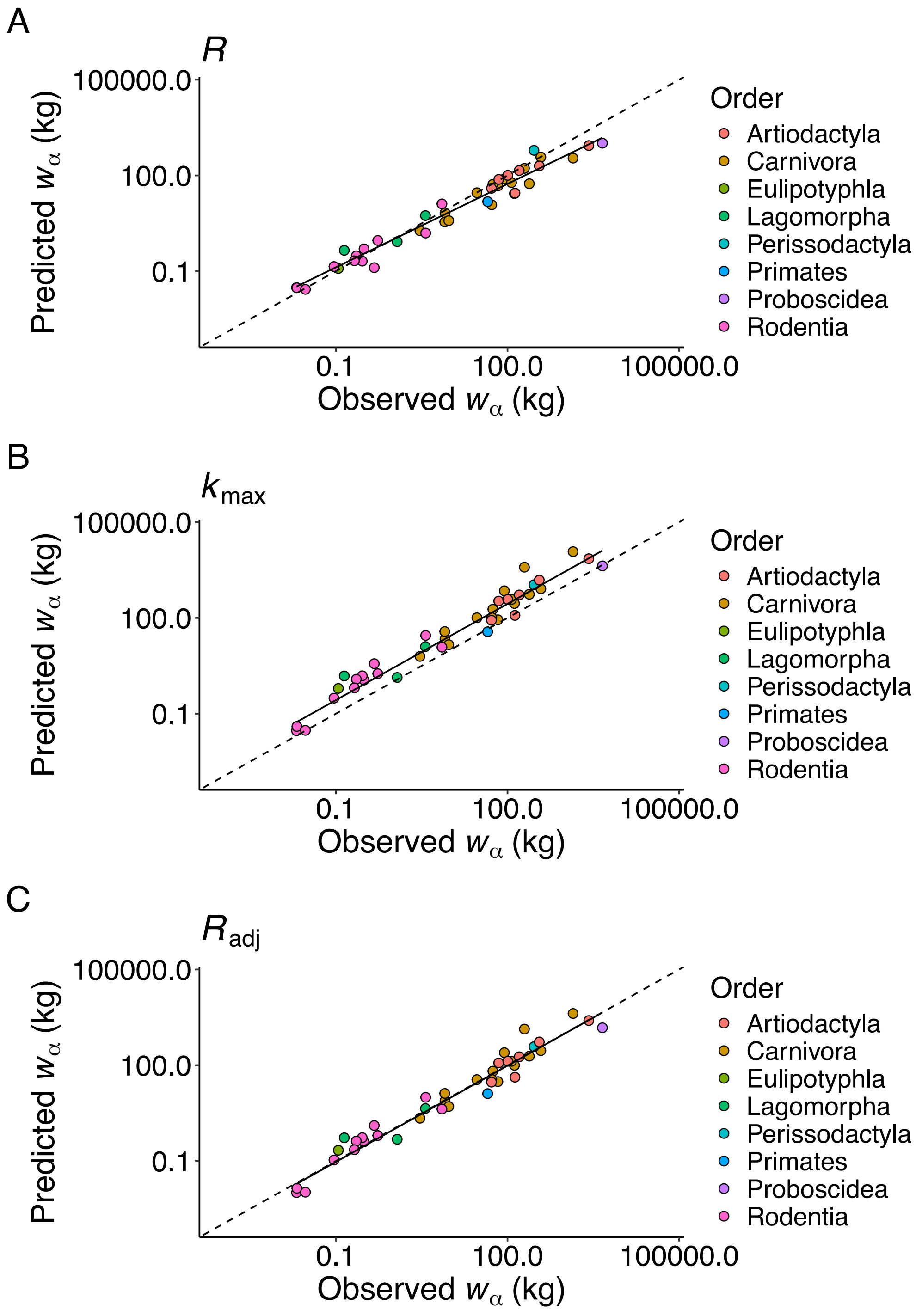


*Figure 3. Adjusting the reproduction rate. PGLS log-log regressions of predicted adult body mass on observed mass using the Demography dataset. (A) Reproduction rate:* $\lambda = 0.86, \kappa = 1.12, \delta = 3.00; r^2 = 0.94, F_{1,39} = 618.6,\ P < 2.20 \times 10^{-16}; y = -0.01 + 0.87x$. *(B) Maximum growth rate:* $\lambda = 0.93, \kappa = 1.09, \delta = 2.92; r^2 = 0.94, F_{1,39} = 609.1,\ P < 2.20 \times 10^{-16}; y = 0.43 + 1.00x$. *(C) Adjusted reproduction rate:* $\lambda = 0.93, \kappa = 1.09, \delta = 2.92; r^2 = 0.94, F_{1,39} = 609.1,\ P < 2.20 \times 10^{-16}; y = -0.02 + 1.00x$. *Dashed lines indicates* $y = x$.


1 Charnov, E. L. & Berrigan, D. Dimensionless numbers and life history evolution: age of maturity versus the adult lifespan. *Evol Ecol* **4**, 273-275 (1990).

2 Charnov, E. L. *Life History Invariants: Some Explorations of Symmetry in Evolutionary Ecology*. (Oxford University Press, 1993).

3 Sibly, R. M., Barker, D., Denham, M. C., Hone, J. & Pagel, M. On the regulation of populations of mammals, birds, fish, and insects. *Science* **309**, 607-610 (2005). https://doi.org/DOI: 10.1126/science.1110760

4 Sibly, R. M., Barker, D., Hone, J. & Pagel, M. On the stability of populations of mammals, birds, fish and insects. *Ecology Letters* **10**, 970-976 (2007).

5 Brook, B. W. & Bradshaw, C. J. Strength of evidence for density dependence in abundance time series of 1198 species. *Ecology* **87**, 1445-1451 (2006).

6 Sinclair, A. R. E. in *Frontiers of population ecology* (eds R.B. Floyd, A.W. Sheppard, & P.J. De Barro) 127-154 (CSIRO Publishing, 1996).

7 Bonenfant, C. *et al.* Empirical evidence of density-dependence in populations of large herbivores. *Advances in Ecological Research* **41**, 313-357 (2009). https://doi.org/10.1016/s0065-2504(09)00405-x

8 Sinclair, A. in *Ecological concepts* (ed J.M. Cherrett) 197-241 (Blackwell, 1989).

9 Gaillard, J.-M., Festa-Bianchet, M. & Yoccoz, N. G. Population dynamics of large herbivores: variable recruitment with constant adult survival. *Trends in Ecology & Evolution* **13**, 58-63 (1998). https://doi.org/10.1016/S0169-5347(97)01237-8

10 Fowler, C. W. in *Current Mammalogy* (ed H. H. Genoways) Ch. 10, 401-441 (Springer, 1987).

11 Fowler, C. W. Density dependence as related to life history strategy. *Ecology* **62**, 602-610 (1981). https://doi.org/https://doi.org/10.2307/1937727

12 Charlesworth, B. *Evolution in Age-Structured Populations*. 2nd edn, (Cambridge University Press, 1994).

13 Sutherland, W. J., Grafen, A. & Harvey, P. H. Life-history correlations and demography. *Nature* **320**, 88-88 (1986). https://doi.org/DOI 10.1038/320088a0

14 Charnov, E. L. Trade-off-invariant rules for evolutionarily stable life histories. *Nature* **387**, 393-394 (1997). https://doi.org/DOI 10.1038/387393a0

15 Charnov, E. L. Evolution of life-history variation among female mammals. *Proceedings of the National Academy of Sciences of the United States of America* **88**, 1134-1137 (1991). https://doi.org/DOI 10.1073/pnas.88.4.1134

16 Tidière, M. *et al.* Comparative analyses of longevity and senescence reveal variable survival benefits of living in zoos across mammals. *Scientific Reports* **6**, 36361 (2016). https://doi.org/10.1038/srep36361

17 Ricklefs, R. E. & Cadena, C. D. Lifespan is unrelated to investment in reproduction in populations of mammals and birds in captivity. *Ecology letters* **10**, 867-872 (2007).

18 Lynch, H. J., Zeigler, S., Wells, L., Ballou, J. D. & Fagan, W. F. Survivorship patterns in captive mammalian populations: implications for estimating population growth rates. *Ecological Applications* **20**, 2334-2345 (2010).

19 Mylius, S. D. & Diekmann, O. On evolutionarily stable life histories, optimization and the need to be specific about density dependence. *Oikos* **74**, 218-224 (1995). https://doi.org/10.2307/3545651

20 Kozłowski, J., Konarzewski, M. & Gawelczyk, A. in *Macroecology: Concepts and Consequences* (eds T.M. Blackburn & K.J. Gaston) Ch. 16, 299-320 (Blackwell Publishing, 2003).

21 Kozłowski, J. & Wiegert, R. G. Optimal age and size at maturity in annuals and perennials with determinate growth. *Evol Ecol* **1**, 231-244 (1987). https://doi.org/Doi 10.1007/Bf02067553

22 Charnov, E. L. Evolution of mammal life histories. *Evolutionary Ecology Research* **3**, 521-535 (2001).
23 Ronget, V. *et al.* Causes and consequences of variation in offspring body mass: meta-analyses in birds and mammals. *Biological Reviews* **93**, 1-27 (2018). https://doi.org/https://doi.org/10.1111/brv.12329
24 Charnov, E. L. Mammal life-history evolution with size-dependent mortality. *Evolutionary Ecology Research* **7**, 795-799 (2005).
25 Charnov, E. L., Warne, R. & Moses, M. Lifetime reproductive effort. *The American Naturalist* **170**, E129-E142 (2007). https://doi.org/https://doi.org/10.1086/522840
26 Ginther, S. C., Cameron, H., White, C. R. & Marshall, D. J. Metabolic loads and the costs of metazoan reproduction. *Science* **384**, 763-767 (2024). https://doi.org/doi:10.1126/science.adk6772
